# Computational structural genomics unravels common folds and predicted functions in the secretome of fungal phytopathogen *Magnaporthe oryzae*

**DOI:** 10.1101/2021.01.25.427855

**Authors:** Kyungyong Seong, Ksenia V Krasileva

## Abstract

*Magnaporthe oryzae* relies on a diverse collection of secreted effector proteins to reprogram the host metabolic and immune responses for the pathogen’s benefit. Characterization of the effectors is thus critical for understanding the biology and host infection mechanisms of this phytopathogen. In rapid, divergent effector evolution, structural information has the potential to illuminate the unknown aspects of effectors that sequence analyses alone cannot reveal. It has recently become feasible to reliably predict the protein structures without depending on homologous templates. In this study, we tested structure modeling on 1854 secreted proteins from *M. oryzae* and evaluated success and obstacles involved in effector structure prediction. With sensitive homology search and structure-based clustering, we defined both distantly related homologous groups and structurally related analogous groups. With this dataset, we propose sequence-unrelated, structurally similar effectors are a common theme in *M. oryzae* and possibly in other phytopathogens. We incorporated the predicted models for structure-based annotations, molecular docking and evolutionary analyses to demonstrate how the predicted structures can deepen our understanding of effector biology. We also provide new experimentally testable structure-derived hypotheses of effector functions. Collectively, we propose that computational structural genomic approaches can now be an integral part of studying effector biology and provide valuable resources that were inaccessible before the advent of reliable, machine learning-based structure prediction.

## Introduction

*Magnaporthe oryzae* is one of the most destructive fungal pathogens that threaten agriculture and food security worldwide (1, 2). The pathogen has adapted to important Poaceae crops, such as rice, millet and barley, and causes destructive blast disease typically accompanying yield losses (3). Domestication and cultivation of crops provides continuous opportunities for the pathogen to develop new virulences (4). A wheat-infecting strain was initially reported in Paraná state of Brazil in 1985 (5). A severe outbreak by related strains swept wheat fields in Bangladesh in 2016 (6).

*M. oryzae* is hemibiotrophic. The pathogen first establishes a biotrophic relationship with its hosts, suppressing their innate immunity and exploiting living host cells (7). In the transition to the necrotrophic phase, *M. oryzae* mediates host cell death and harvests nutrients from the dead tissues. The success of the infection cycle depends on a diverse set of effectors *M. oryzae* secretes during its interaction with the host plants. Plant innate immune systems are sophisticated enough to perceive the presence of invading pathogens and their activity (8). Specifically, extracellular cell-surface receptors recognize conserved pathogen-associated molecular patterns (PAMPs), such as chitin, and mediate PAMP-triggered immunity (PTI). Inside the plant cell, nucleotide-binding and leucine-rich repeat receptors can detect cytoplasmic effectors and induce effector-triggered immunity. The recognition of effectors by their cognate receptors denote their avirulence, but to the pathogen, this may be directly linked to the deprivation of its opportunity to survive. *M. oryzae*, therefore, has rapidly evolved to evade and suppress immune responses (9).

Signatures of rapid effector evolution remain in the pathogens’ genomes. A genomic study identified a bipartite genome architecture where the known avirulence genes are primarily located in repeat-rich, gene-sparse regions with rapid changes (10). Evolutionary analyses uncovered lineage-specific presence and absence variations of avirulence effectors and the mechanisms of gene flow and recombination that contribute to diversifying the effectorome (3, 11–14). Extensive molecular characterization has been applied to unravel the biological roles and molecular functions of the virulence factors; however, our present understanding of effector biology is fragmented, and most of the effectors remain to be characterized.

Structural biology has the potential to illuminate effector biology, as the structural commonality may be an important aspect of effector evolution typically accompanying sequence divergence (15). Avirulent effectors Avr1-CO39, Avr-Pia and AvrPiz-t from *M. oryzae* and ToxB from the wheat pathogen *Pyrenophora tritici-repentis* are unrelated by their sequences; however, they share a common β-sandwich fold and are together classified as MAX (*<u>M</u> agnaporthe* <u>A</u>vrs and <u>T</u>oxB-like) effectors (16). Similarly, in phytopathogenic *Leptosphaeria maculans*, AvrLm3, AvrLm4-7 and AvrLm5-9 are structural analogs belonging to LARS (*<u>L</u> eptosphaeria* <u>a</u>virulence-<u>s</u>uppressing) effectors (17, 18). In powdery mildews, such as *Blumeria graminis*, RALPH (<u>R</u>N<u>a</u>se-like <u>p</u>roteins expressed in <u>h</u>austoria) effectors, that commonly adopt the structural folds of RNase despite highly divergent sequences, represent a significant portion of the effectorome (19–21). The advantages of protein structures in understanding effector biology, therefore, make it particularly more valuable to incorporate the principle of structural genomics, an approach that aims to determine the structures for all proteins by experimental and prediction methods (22).

The application of structural genomics to effectors has, however, remained limited. Experimental determination of all effector structures would require immense time and community effort. Prediction is an attractive alternative; nonetheless, the accurate prediction has, until recently, been predominantly derived from homology modeling unsuitable for effectors that typically lack homologous templates. In the Critical Assessment of Structure Prediction (CASP) 13 in 2018, the success of AlphaFold opened a new path around this impediment (23, 24). By inferring co-evolutionary features from a collection of homologs and integrating deep learning, the structure of a protein could be reliably predicted in the absence of homologous templates. Soon, powerful tools like TrRosetta became available for research, allowing us to examine the feasibility of applying the computational structural genomics approaches to understand the secretome of *M. oryzae* (25). In the present study, we modeled the structures of 1854 secreted proteins with template-based modeling by I-TASSER and deep learning-incorporated free-modeling by TrRosetta (26). Using the structural information, we provide new hypotheses that can inform future experimental work and bridge gaps in our understanding of M. oryzae effector biology.

## Results

### Benchmarking on *Magnaporthe* proteins with solved structures suggests the folds of many effectors can be predicted without homologous templates

Prior to the structure prediction on *M. oryzae* secretome, we assessed the performance of the prediction tools on representative 9 secreted and 15 non-secreted *Magnaporthe* proteins available in the Protein Data Bank (PDB) (Table S1, Fig. S1) (27). We predicted their structures with TrRosetta and I-TASSER and superposed the computational models against the experimental ones with TM-align (28). In this benchmarking, non-secreted proteins with functional annotations were easy targets for both methods, given the high precision of the predicted structures (Fig. 1*A*). TrRosetta produced structures with TM-scores > 0.5 for known avirulence proteins AvrPiz-t (2LW6), Avr-Pia (2N37), Avr-Pib (5Z1V) and Avr-PikD (6G10), while I-TASSER only predicted the Avr-Pia structure (Fig. S2). Such levels of similarity indicate that the predicted and native structures display about the same fold (29). TrRosetta utilizes residues’ coevolutionary information from a multiple sequence alignment (MSA) composed of the target protein and its homologs to infer distances and orientations of residues. However, the MSAs of the avirulence proteins were shallow with the number of homologs ≤ 13 and likely insufficient for such inference (Table S1). Therefore, the prediction results rather suggested that trained machine learning algorithms alone were sufficient to capture the overall folds of these effector structures.

**Figure 1.**
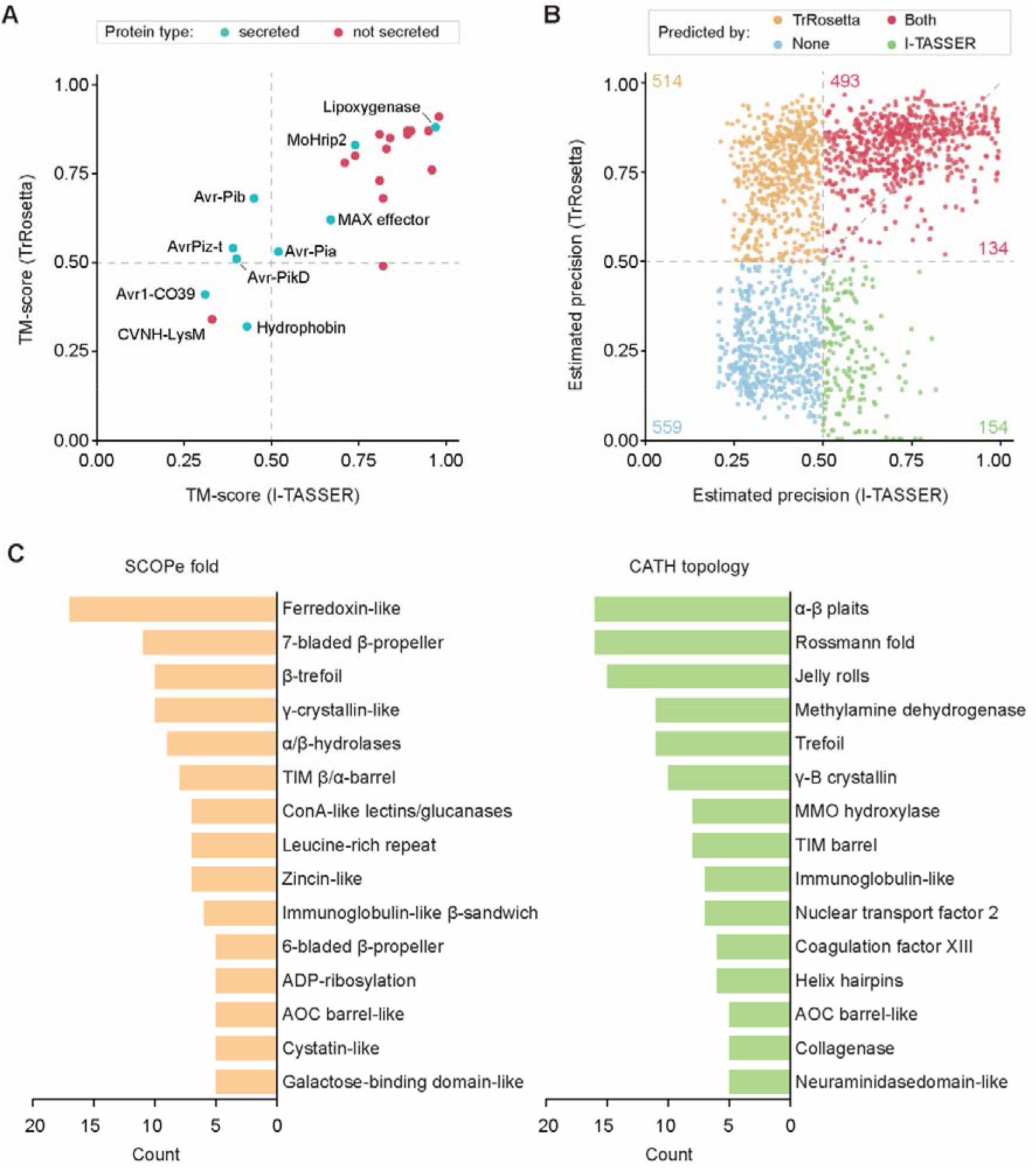
The statistics of protein structure prediction on *Magnaporthe* proteins with solved structures available in the Protein Data Bank and the putative secreted proteins in the *Magnaporthe oryzae* genome. **A**. The precision of the structures predicted by TrRosetta and I-TASSER for the 24 *Magnaporthe* proteins available in the Protein Data Bank. For homology modeling, templates with ≥ 70% sequence identity were excluded. The TM-scores were obtained by superposing the computational models against the experimental structures with TM-align. A TM-score > 0.5 indicates the compared structures display about the same fold. Secreted and non-secreted proteins are noted in different colors. **B**. The estimated precision of the structures predicted by TrRosetta and I-TASSER. The average probability of the top L predicted long+medium range contacts (|i-j|>12) and the mean estimated TM-score were used as the expected precision for TrRosetta and I-TASSER, respectively. These values are reported to well correlate with the actual precision. The number of proteins belonging to each section is indicated in the plot. **C**. The structure-based classification of 527 uncharacterized proteins without functional annotations. The 527 uncharacterized proteins predicted with the estimated precision > 0.5 were assigned to SCOPe and CATH categories with RUPEE. The TM-score cut-off > 0.5 was required for the classification. The top 15 hits are reported.

Despite sharing common folds with AvrPiz-t and Avr-Pia, Avr1-CO39 (5ZNG) was a challenging target for both methods with the noticeable discrepancy in secondary structures (Fig. S2) (30). Also, secreted hydrophobins (2N4O) and the CVNH-LysM dual-domain (2L9Y) failed to be predicted with TM-scores > 0.5. For both proteins, a sufficient number of homologs existed for TrRosetta, and I-TASSER identified homologous templates (2NBH and 5JCE, respectively). Possibly, a high-level of structural or sequence diversification among homologs would explain the predicted topologies dissimilar to their native structures for hydrophobins (31). An atypical architecture of the bisected CVNH domain could be problematic, as the inserted LysM domain was correctly modeled (Fig. S3) (32). Collectively, the benchmarking results suggested the structural folds of many effectors could be predicted without homologous templates, although the sequences of a target protein and the evolution of its homologs would have impacts on the prediction outcomes.

### Structural modeling resolves a large subset of secreted protein structures

We next modeled 1854 putative secreted proteins encoded in the *M. oryzae* genome. The estimated precision TrRosetta and I-TASSER report for a predicted structure well-correlates with the actual precision (25, 33). For TrRosetta, we also obtained a previously reported correlation coefficient of 0.84 on our benchmarking data (Fig. S1). We, therefore, counted on this estimate to determine the reliability of the structures. TrRosetta and I-TASSER predicted 627 structures in common with expected precision > 0.5 (Fig. 1*B*, Table S2). Among them, 493 TrRosetta models displayed equal or higher expected precision. Each tool predicted additional 514 and 154 high-confidence structures, respectively (Fig. 1*B*). Collectively, 70% of the secreted proteins were modeled by at least one of the methods with expected precision > 0.5. The lack of homologous templates explains the relatively poor performance of I-TASSER. Many models exclusively modeled by I-TASSER included very short or highly disordered proteins that may not be trustworthy (Table S2; Fig. S4). Among the 559 proteins not predicted, 340 (61%) and 477 (85%) proteins were smaller than 150 and 250 amino acids, respectively. Of the 477 proteins, 335 proteins displayed the proportion of disordered residues < 0.25, indicating they are likely foldable. However, most of the proteins lacked detected homologs for the co-evolutionary inference. Nonetheless, 527 out of 1062 uncharacterized proteins without PFAM domains or with the domain of unknown functions (DUFs) had predicted structures.

We classified the 527 uncharacterized proteins by searching for similar structural folds and topologies in the SCOPe and CATH databases with RUPEE (Fig. 1*C*; Table S3) (34–36). The classification identified enzymes of diverse functions. These include hydrolytic enzymes adopting the α/β-hydrolase and Rossmann fold; glycosidases displaying the TIM (β/α) barrel structure; glucanases belonging to the concanavalin A-like lectins/glucanase and jelly roll structures; and metallopeptidase of the zincin-like and collagenase fold. Some proteins predicted to have the 6-bladed β-propeller and neuraminidase or 7-bladed β-propeller and methylamine dehydrogenase folds were also correlated with enzymatic activities. Collectively, the classification indicated that a subset of the unknown *M. oryzae* secretome contains divergently or rapidly evolving enzymes of diverse functions. Other structural folds and topologies were also present, which may supplement the infection mechanisms of *M. oryzae*. Nonetheless, the classification is not representative of the entire uncharacterized effecterome. Such structural classification requires not only predicted structures of good quality but also the existence of similar structures and their pre-assignment into proper categories. As the solved structures of effectors are limited, novel structural classes of effectors could not be represented in this process.

### Predicted structures for known effectors can be used to infer their possible biological roles

*M. oryzae* encodes a diverse set of cloned effectors, the structures and functions of which are currently unknown (Table S4). We examined whether TrRosetta and I-TASSER modeled any of these effector structures. Some of the effectors have homologous structures from other organisms in the PDB easily identifiable with BLASTP. All of these effectors, except for MoCDIP12 only partially homologous to AVR-Pik (6FUD), were predicted, and in most cases, the TrRosetta models displayed higher expected precision (Fig. S5). In structural comparisons, all but SLP1 significantly resembled their homologous structures, supporting the quality of the prediction (Fig. S6). For SLP1, the LysM domain was well superimposed, but the overall topology was imprecise as the linkers were incorrectly predicted (Fig. S6*H*). Among the effectors without homologous structures, TrRosetta could generate models with high expected precision for MoNIS1, Avr-Pi54, EMP1, MoCDIP1, MoCDIP5, MoCDIP10, Avr-Pita1 and BAS4 with which reliable downstream analyses can be conducted (Fig. S7*A*-*H*). With lower estimated precision, the structures for PWL, BAS3 and MoCDIP3 were also predicted (Fig. S7*I*-*K*).

We searched for structural matches of the effectors without homologous templates to reveal their putative biological roles. Although our PFAM search did not uncover any conserved domains, structural comparability to pectin lyases and metallopeptidases existed for MoCDIP1 and MoCDIP5, suggesting these cell-death inducing proteins may be enzymes (Fig. 2*A* and 2*B*) (37). Nonetheless, the highly conserved residues found in pectin lyases (1IB4) were absent in MoCDIP1, despite the significant structural resemblance (Fig. 2*D*) (38). Similarly for MoCDIP5, only the strictly conserved metallopeptidase HEXXH motif was structurally aligned to the metzincins motif (HEXXHXXGXXH) of the match, collectively indicating that the mode of catalytic mechanisms likely differs for these MoCDIPs (Fig. 2*E*) (39). At the C-terminal region of MoCDIP10, the immunoglobulin-like β-sandwich fold was newly detected (Fig. 2*C*) (40). The DNA-binding TIG domain of human calmodulin-binding transcription activator 1 (2CXK) and other protein-binding domains appeared as similar structures (Fig. 2*F*). Possibly, the β-sandwich fold aids binding to host targets which the ferritin-like domain then acts on. Such structure-based inference of effectors’ putative biological roles can supplement the sequence-based detection. Alternatively, hidden Markov models from the SUPERFAMILY and CATH-Gene3D databases constructed with known structures are necessary to detect some of these domains PFAM cannot annotate (41, 42). Nevertheless, ambiguity remains for some effectors, as they are analogous to structures of diverse functions. For instance, Avr-Pi54 resembles a carbohydrate-binding module (2W47), a hydrolase inhibitor (4DZG) and nucleoplasmin (2VTX), which could not point to a specific role of this effector (Fig. S8).

**Figure 2.**
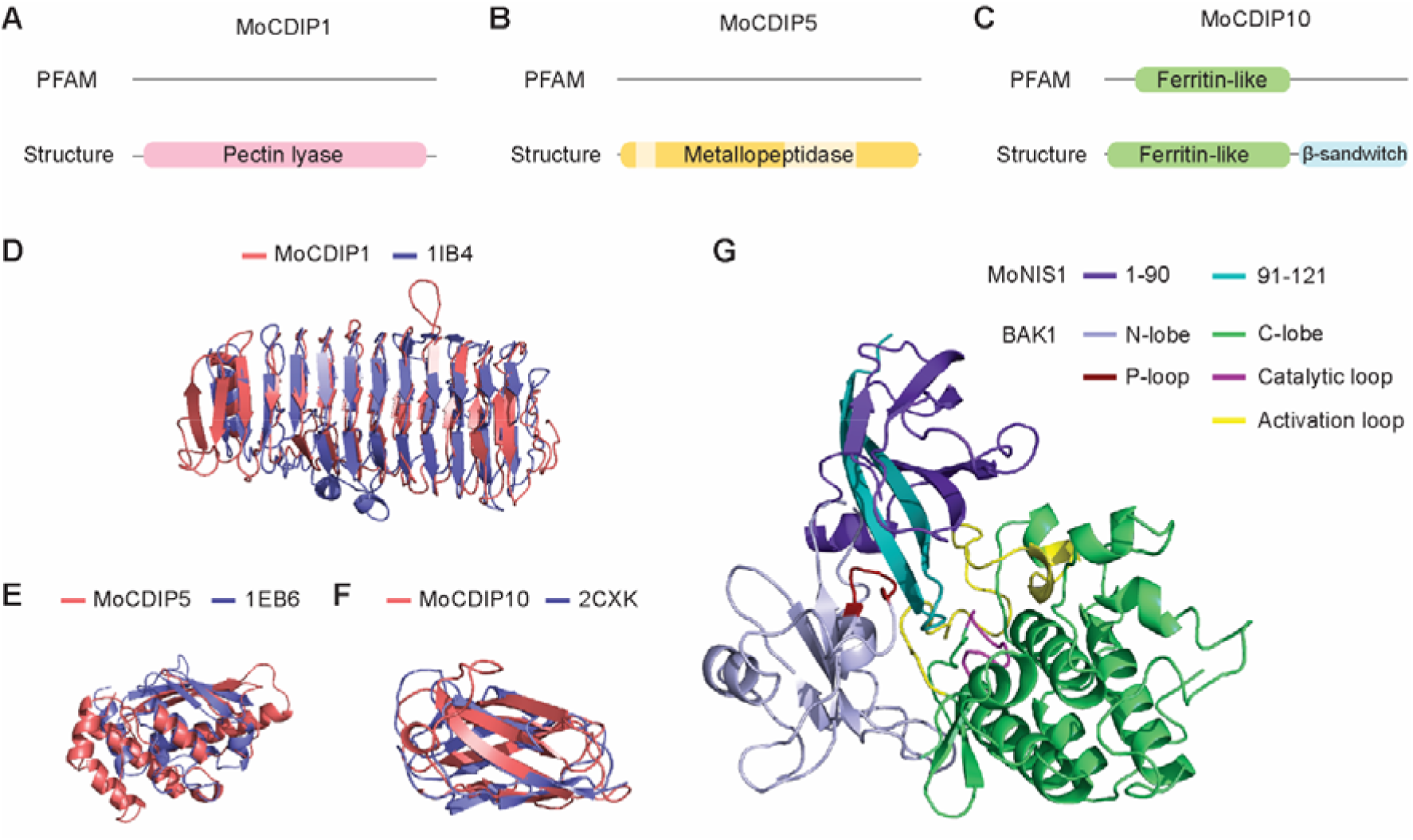
Structure-based inference of effector’s biological roles. **A-C**. The PFAM and structure-based annotations of MoCDIP1, 5 and 10. The PFAM domain search and structural similarity search with RUPEE against the SCOPe and CATH databases were used for the annotation. **B**. The structurally unaligned region of MoCDIP5 is indicated in lighter yellow. **D**-**F**. The structural superposition of MoCDIP1, 5 and 10 and their significant matches. The normalized TM-scores, as a measure of similarity, were 0.75 and 0.74 for MoCDIP1 and 1IB4, 0.52 and 0.61 for MoCDIP5 and 1EB6, and 0.64 and 0.63 for MoCDIP10 and 2CXK. **G**. The docking of MoNIS1 to BAK1 from *Arabidopsis thaliana* (3ULZ). The C-terminal 30 amino acids of MoNIS1 are indicated in cyan. The P-loop, catalytic loop and activation loop are colored in red, yellow and purple.

TrRosseta predicted an immunoglobulin-like fold for MoNIS1. Despite the expected precision of 0.97, MoNIS1 only revealed high similarity to biologically irrelevant matches, such as sterol binding protein (2HKA) and a collagenase domain (2Y72). Unexpectedly, a synthetic monobody (5G15) appeared as a similar structure, as we traced analogs that may contain the predicted MoNIS1 structure (43). This monobody was designed to bind to the oncogenic human protein kinase AurA to allosterically control its kinase activity and co-crystallized with the receptor (Fig. S9*A*). Analogously, MoNIS1 interacts with the cytoplasmic kinase of the cell surface receptor BAK1 in the host, preventing PTI (44). The available structure of BAK1 (3ULZ) displayed noticeable similarity to AurA, possibly suggesting that the interaction between the monobody and AurA might be comparable to that between MoNIS1 and BAK1 (Fig. S9*B* and S9*C*). We, therefore, attempted to dock MoNIS1 into BAK1 with ZDOCK (45). The best docking simulation suggested that MoNIS1 may compete for the substrate-binding site, thereby interfering with the access to the P-loop, activation loop and catalytic loop of BAK1 (Fig. 2*G*) (46). This interaction is mediated by the C-terminal 30 amino acids, the deletion of which was shown to abolish the role of MoNIS1 homolog in *Colletotrichum* spp. (44). Such interaction is rather analogous to the one between BAK1 and AvrPtoB from bacterial pathogen *Pseudomonas syringae* (47). The lack of structural and evolutionary similarity between the effectors points to convergent evolution in the effector mechanism in bacterial and fungal phytopathogens.

### Sensitive similarity search and structural comparisons define secretome classes in *M. oryzae*

Many effectors evolve rapidly and divergently to evade host immune recognition (9). As the effectors substantially deviate from other canonical homologs in sequence, homology detection failure with BLASTP becomes more likely and frequent. In such cases, more sensitive methods or structural comparison would aid distant homology identification. To unravel the interconnection between the secreted proteins in *M. oryzae*, we performed secretome clustering with these methods.

As we first detected sequence-to-sequence homology with BLASTP, 819 proteins found at least another homolog present in the secretome (Fig. 3*A*; Table S5) (48). With the MSA we constructed for each secreted protein and its homologs, we then performed a more sensitive profile-to-sequence similarity search with HHblits, linking additional 186 sequences into homologous groups (49). Profile-to-profile similarity search with HHsearch and structural comparisons with TM-align added 49 and 69 proteins to the clusters, respectively. Eventually, 1123 proteins (60%) had at least another related protein in the secretome, 449 of which did not contain PFAM domains. 731 proteins remained as singletons that lack detectable paralogs and structural analogs in the secretome, and only 118 of them contained functional annotations (Fig. 3*B*). Of the 613 remaining proteins without PFAM annotations, only 86 proteins had predicted structures with the estimated precision ≥ 0.6, which could be used more reliably for subsequent analyses.

**Figure 3.**
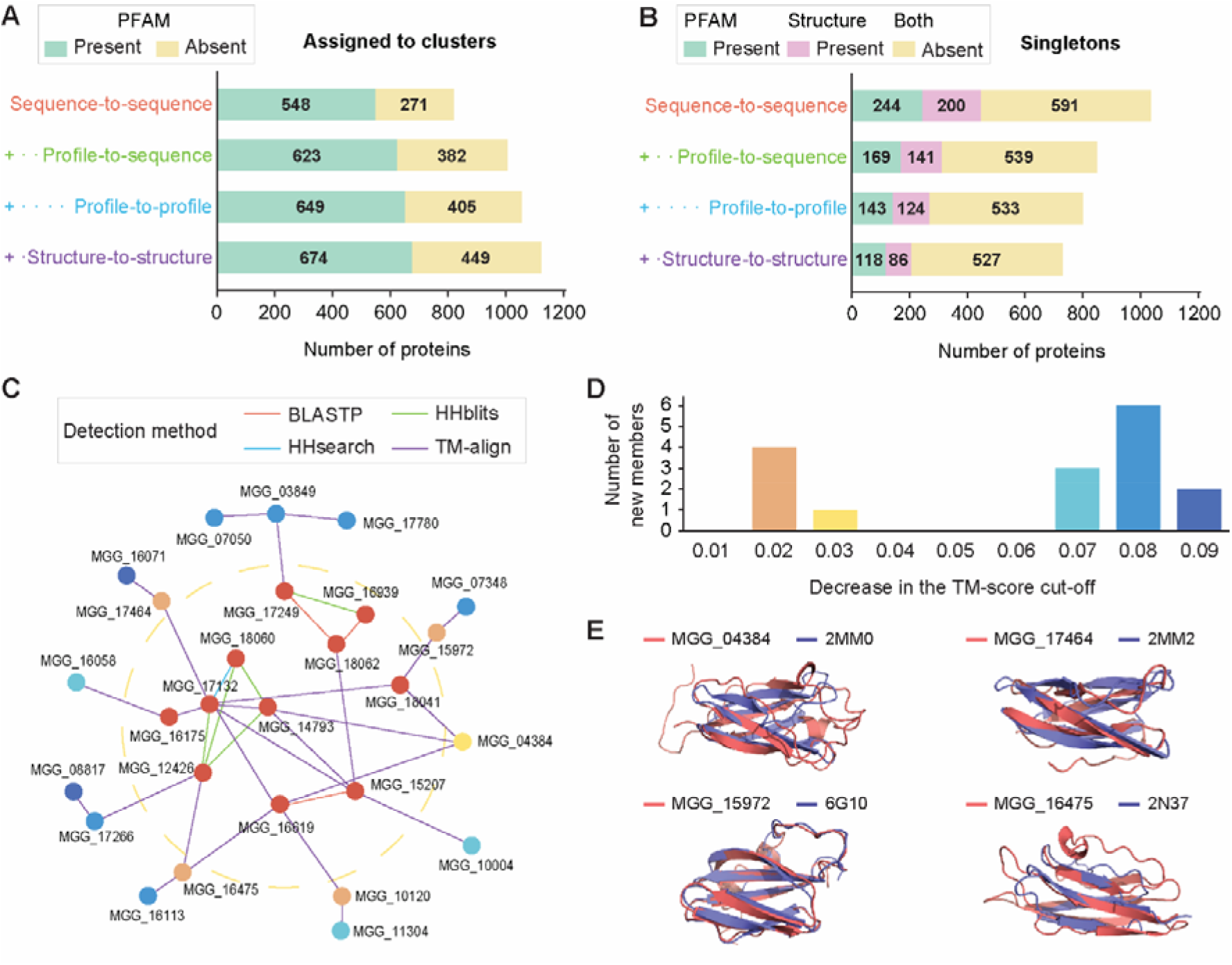
Statistics on secreteome clustering and the MAX effector cluster. **A**-**B**. Four methods were sequentially used to reveal sequence-based homology and structure-based homology or analogy. **A**.The number of proteins found in clusters with at least another homolog or structural analog in the secretome. **B**. The number of singletons without any homologs or analogs in the secretome. The number of proteins with meaningful PFAM domains, excluding domain of unknown functions, or structures predicted with estimated precision ≥ 0.6 are indicated. **C**-**D**. The network graph for MAX effectors and the number of newly retrieved singletons. **C**. Each node and edge represent a protein and similarity that can be detected by the method. The MAX effector cluster with 11 members (Cluster 26) exists inside the yellow dotted ring. The newly retrieved singletons remain outside the ring. **D**. The criteria for the final model selection (> 0.5 for TrRosetta and > 0.55 for I-TASSER) and the significant structural similarity (> 0.5 for both of the compared structures) were relaxed by the TM-score of 0.01. The number of newly retrieved singleton members is indicated. The colors correspond in **C** and **D. E**. The structural superposition of the newly retrieved MAX effector members and their most similar structures available in the Protein Data Bank. The normalized TM-scores, as a measure of similarity, were 0.47 and 0.63 for MGG_04384 and 2MM0; 0.47 and 0.46 for MGG_17464 and 2MM2; 0.81 and 0.89 for MGG_15972 and 6G10; and 0.47 and 0.60 for MGG_16475 and 2N37.

We specifically traced putative MAX effectors, as sensitive homology search and structural comparisons are required to reveal them (16). In the final clusters we generated, a group of 11 uncharacterized proteins, which includes homologs of Avr-Pib (MGG_12426) and AvrPiz-t (MGG_18041), was grouped by sequence or structural similarity in Cluster 26 (Fig. 3*C*; Table S5). In sequence-to-sequence similarity search, these 11 proteins were separated into 6 singletons and two clusters. By the profile-to-sequence homology search, two singletons and three individual clusters formed, which did not display sufficient sequence similarity to each other. However, the structure-to-structure similarity search was able to link them all, eventually placing the Avr-Pib and AvrPiz-t homologs into the single cluster.

Although the sensitive homology search and structural comparisons could define the putative MAX effector cluster, a homolog of Avr-PikD (MGG_15972) remained as a singleton. MGG_15972 had a predicted structure with a relatively high precision of 0.77, yet it slightly missed the criterion of significant structural similarity measured with the TM-score > 0.5 for both compared structures. As this happens for the native MAX effector structures, and some MAX effectors are predicted with lower estimated precision, we reduced the cut-offs for the final model selection and structural similarity comparisons by a TM-score of 0.1 (Fig. 1*A*; Fig. S10). We then examined whether any singletons could be retrieved into this cluster (Fig. 3*D*). By decreasing the cut-off by 0.03, we found five new members, four of which identified *P. tritici-repentis* ToxB (2MM0 and 2MM2), Avr-PikD (6G10) and Avr-Pia (2N37) as their most similar structures in the PDB, respectively (Fig. 3*E*). Additional members appeared only after the cut-off was reduced by 0.07, and by 0.09, 11 new members were retrieved. However, only one of them (MGG_17266) displayed similarity to Avr-Pib (5Z1V) with an average TM-score of 0.45, while the other members found unrelated matches (Table S5). Such outcomes indicate that the sensitivity of the structure-based clustering can be enhanced by relaxing its criteria, yet this likely comes at the cost of precision.

### Secretome clustering and structure-based functional inference lead to new hypotheses for the infection mechanism of *M. oryzae*

Secretome clustering and inferring functions of each cluster facilitate understanding the putative roles of effectors. Especially, the structure-based functional inference can bridge the gap across rapidly diverging sequences, leading to new hypotheses. A recent study demonstrated that the two nuclear effectors, MoHTR1 and MoHTR2, bind to the promoters of immunity-related genes in rice and interfere with their expression (50). In our clustering, MoHTR2 along with Nup3, also shown to block plant immunity, belong to Cluster 25 with 9 other members with distant homology (Table S4; Table S5) (51). The presence of nucleic acid-binding zinc-finger domain (PF12171) in this cluster well-correlates with the actual molecular functions, and the size of the cluster suggests that this mechanism may be important to *M. oryzae*. Host nucleic acids can be also targeted by RNases as demonstrated by RNase-like effectors that significantly constitute the effectorome of *B. graminis*. The structure prediction and comparisons identified a small RNases cluster in the *M. oryzae* secretome (Cluster 74). The predicted structures belong to the T1 family with similarity to the *B. graminis* RNase-like effector (6FMB), pointing to the possible existence of a common mechanism in the two distant phytopathogens (Fig. S11).

We formulate another hypothesis on possible carbohydrate-binding proteins. TrRosetta predicted with high estimated precision the structures for necrosis-inducing factor domain (PF14856), also known as the Hce2 (<u>h</u>omologs of *<u>C</u>. fulvum* <u>E</u>cp2 effectors) domain (52). The predicted model identified glycan-binding protein Y3 isolated from fungus *Coprinus comatus* (5V6J) as the closest analog and was linked to 7 other sequence-unrelated members by structural analogy in Cluster 33 (Table S5). The structural similarity to Y3 is limited to the fold level, as the TM-score was only about 0.57. However, as the previous study identified co-occurrence of the Hce2 domain in the proteins containing other lectin-binding LysM domains, chitin-binding modules and chitinase, it may be possible that the Hec2 domain is involved in their common biological goal of glycan-binding and processing (53).

### Sequence-unrelated structural analogs may be a common theme in effector evolution

The MAX effector cluster is the best representative of the sequence-unrelated, structurally similar effectors in *M. oryzae*. Other such structurally analogous clusters may exist, yet examining them has been elusive as the solved structures are limited. With the structure-based clustering, we could capture the presence of sequence-irrelevant, structural common themes (Table S5). An example is BAS4 which is highly expressed at the invasive hyphae and participates in the transition from biotrophy to necrotrophy (54, 55). BAS4 exists in cluster 17 with 14 other members initially divided into five distinct groups by sequence (Fig. 4*A*). However, the predicted structures disclosed a common ferredoxin-like fold and alpha-beta plait topology, linking the groups into a single cluster (Fig. 4*B*). This topology includes proteins of diverse functions, and the predicted structures broadly displayed similarity to them, including a virally encoded killer toxin (4GVB), a metal-binding protein (2GA7) and a chaperone protein (3OFH) (Fig. S12). Despite the predicted structures with relatively high estimated precision, their putative functions, therefore, could not be inferred. Interestingly, the CATH classification assigns LARS effectors, including AvrLm4-7 (4FPR), to the alpha-beta plait topology (17). In comparison to the members in Cluster 17, AvrLm4-7 is larger and displays a difference in the secondary structure topology. Nevertheless, the structural superposition illustrated that MGG_01064 is roughly contained in AvrLm4-7, possibly suggesting that unknown virulence functions associated with this topology may be important for phytopathogens (Fig. 4*C*).

**Figure 4.**
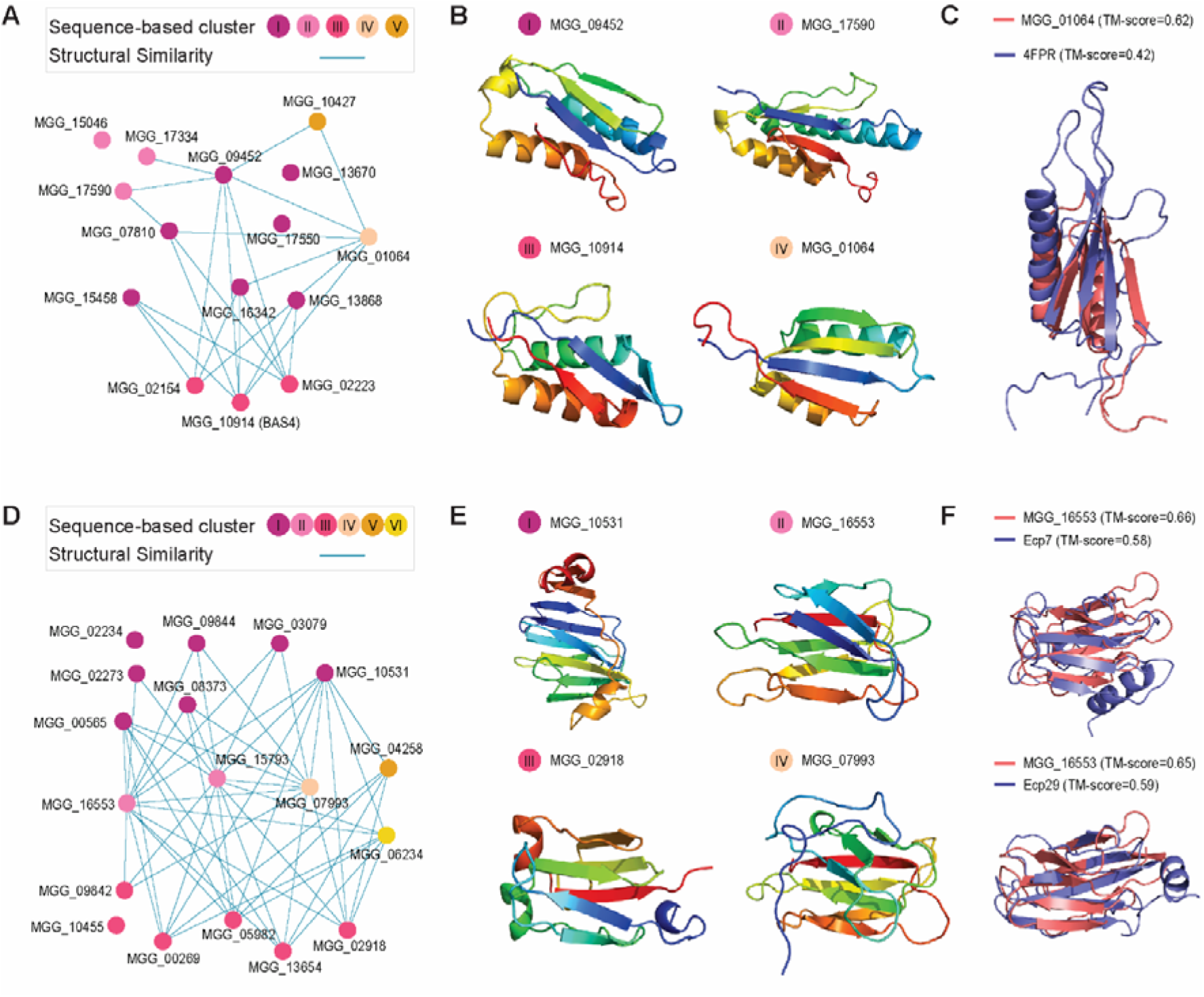
Large clusters of sequence-unrelated structurally similar effectors and the appearance of structural analogs in other phytopathogens. **A** and **D**. The network graph for Cluster 17 and 14, respectively. The node represents a member in the cluster and is colored according to its final membership based on sequence-based homology search. The edge indicates detectable structural similarity. If such similarity is present between two proteins with different membership, their nodes were connected. **B** and **E**. Representative structures for Cluster 17 and 14. The structures predicted with the highest expected precision were selected from each sequence-based cluster. **C**. The structural superposition between the predicted MGG_01064 and LARS effector protein 4FPR. The TM-score, as a measure of similarity, normalized for each structure is indicated in the parentheses. **F**. The structural superposition between the MGG_16533 model and Ecp7 as well as the C-terminal region of Ecp29 (158-266) predicted by TrRosetta.

Cluster 14 also demonstrates sequence-unrelated, structurally similar putative effectors. The 18 members are divergent, and the sequence similarity searches only revealed three homologous groups and three singletons (Fig. 4*D*). However, the members commonly share the γ-crystallin-like fold and are thus structurally related (Fig. 4*E*). This superfamily includes yeast killer toxins (1WKT) and chitin-binding protein secreted from bacterial species *Streptomyces tendae* (1G6E) (Fig. S13). The predicted structures also display significant similarity to other γ-crystallin-like structures with biologically irrelevant functions, such as mammalian eye lens proteins (1A7H). Although the functional inference is not feasible, this group may play an important role in pathogenesis. In *M. oryzae*, lineage-specific presence and absence variations of MGG_16553 and MGG_15793 were detected (56). Additional studies emphasized the exclusive appearance of a *Fusarium pseudograminearum* homolog (FPSE_05488) in cereal-infecting fungi, and a high expression of a *Colletotrichum graminicola* homolog (GLRG_11440) during the transition to biotrophy (57, 58). It was also suggested that four extracellular effectors, Ecp4, Ecp7, Ecp29 and Ecp30, from *Cladosporium fulvum* would adopt a β/γ-crystallin fold (59). Consistently, the TrRosetta server generated reliable structures with the expected fold (Fig. S14). These effectors indeed displayed noticeable similarity to the members in Cluster 14, collectively indicating that these structural analogs may play a significant role in not only *M. oryzae* but also other phytopathogens (Fig. 4*F*).

The structural analogy also appeared for known virulence factors, such as MoHrip1 in Cluster 13 and MoHrip2 in Cluster 44 (Table S4; Table S5). Such trends support that the theme of sequence-unrelated, structurally similar effectors may not be limited to MAX effectors but are likely more common for others that may play an important role in host infection and colonization.

### The ADP-ribosylation fold may be important in *M. oryzae* pathogenicity and evolution

Understanding how *M. oryzae* modulates and interferes with host cellular activities remains vital. A possible mechanism by which the pathogen accomplishes this could be via ADP-ribosylation, the transfer of an ADP-ribose moiety to the host targets in the NAD^+^-dependent manner (Fig. 1*C*). ADP-ribosylation in plant pathogenesis is well represented with type III effectors HopU1 (3U0J) and HopF2 (AvrPphF; 1S21) from *P. syringae* (60–62). The putative ADP-ribose transferases (ARTs) in *M. oryzae*, however, belong to a structurally different class and are associated with the heat-labile enterotoxin alpha chain (PF01375). Although this domain is typically correlated with human diseases mediated by bacteria, such as *Escherichia coli* and *Vibrio cholerae*, homologous Scabin toxin (5EWY) encoded by bacterial phytopathogen *Streptomyces scabies* demonstrated its role in plant pathogenesis (Fig. S15) (63).

Cluster 8 includes 27 members, only four of which contain the heat-labile enterotoxin alpha chain (Table S5). Among the members, ten possessed structures predicted with high estimated precision > 0.75 and the ADP-ribosylation fold. All of these proteins, except for one, formed the largest homologous group that can be simply revealed with BLASTP despite sequence divergence; therefore, we designate them as core ARTs. To elucidate how the core members evolve, we employed Shannon’s entropy on the ten proteins, as it robustly captures sequence diversity (64, 65). Well-conserved residues measured with entropy ≤ 1 contained some known catalytic residues (R-ST-E) and residues around them (Fig. 5*A*) (66, 67). These residues primarily appeared in proximity in the 3D structure, implying their functional importance (Fig. 5*B*). We also superimposed the core members’ structures and Scabin with mTM-align (68). The superposition revealed a highly similar structural core, containing the putative substrate-binding sites and the catalytic sites despite sequence diversification between the paralogs that the entropy distribution indicated (Fig. 5*C*). Consistently, the protein-ligand docking simulation by EDock located an NAD^+^ molecule into this region (Fig. S16) (69). Conversely, for other residues, sequence variations correlated with structural deviations. Such evolutionary patterns possibly suggest that these putative core ARTs have diversified for different host targets and evolved divergently in co-evolution against plant immunity while maintaining their core structures and functions.

**Figure 5.**
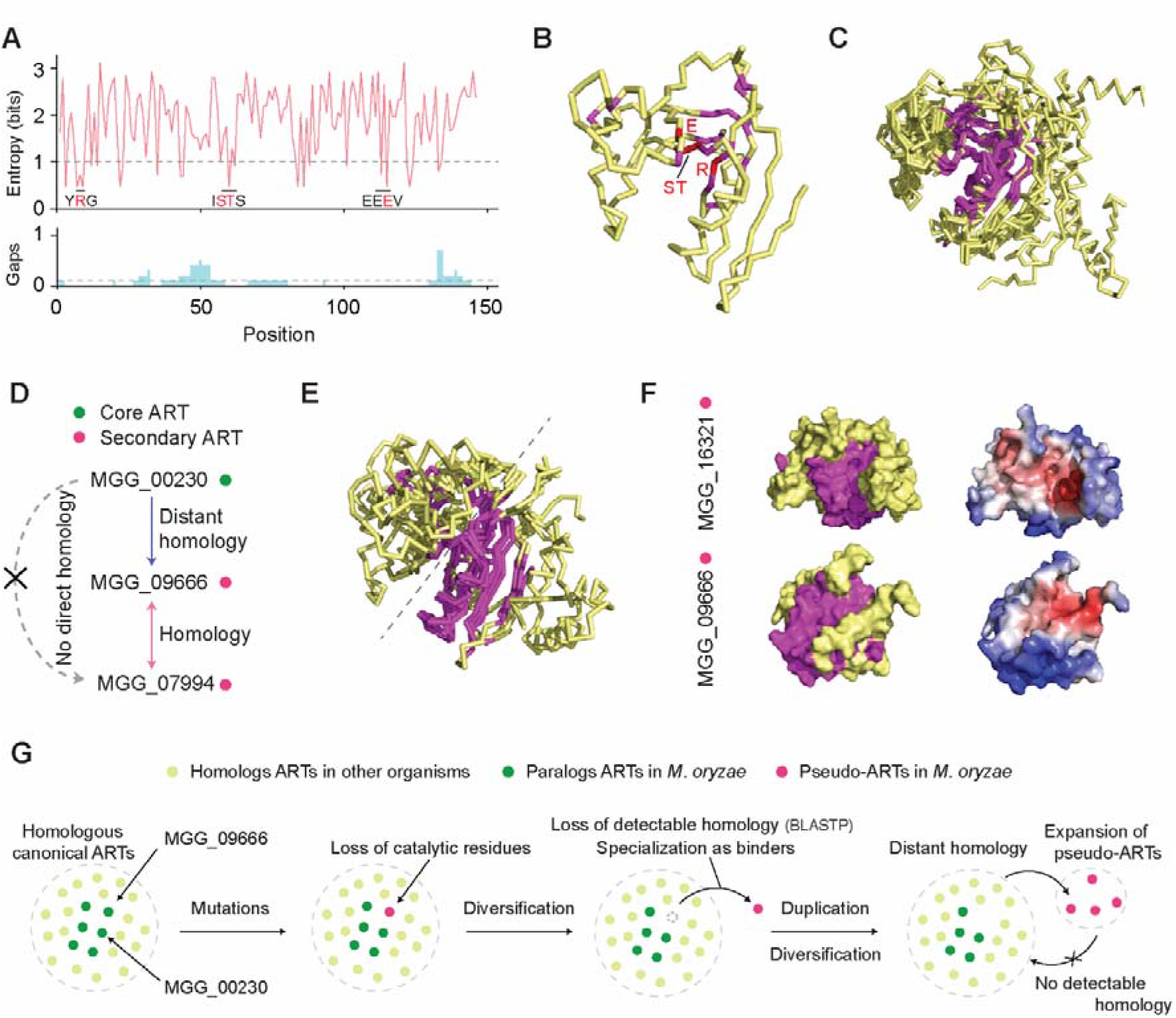
The evolution of putative ADP-ribose transferases. **A**. The entropy plot for the core putative ADP-ribose transferases (ARTs) with structures predicted with estimated precision > 0.75 and the ADP-ribosylation fold. The ten sequences were aligned with MAFFT, and Shannon’s entropy was calculated for the columns that contain MGG_00230 sequences. The gap was ignored in the entropy calculation, but the proportion of gap characters are indicated below the entropy plot. The cut-off for conserved residues was entropy ≤ 1 and the proportion of gap < 0.1. The known catalytic residues (red) and residues around them (black) are indicated in the entropy plot. **B**. The ribbon structure of MGG_00230 with annotated conserved and catalytic residues in magnata and red, respectively. **C**. The structural superposition of the ten core ARTs and Scabin toxin generated with mTM-align. The structural core measured with the maximum pairwise residue distance < 4Å is indicated in magnata. **D**. The homology relationship between the core and secondary ARTs in Cluster 8. The arrow indicates the direction of homology. For instance, MGG_00230 → MGG_09666 denotes that homology to MGG_09666 is detectable when MGG_00230 is used as a query. **E**. The structural superposition of the four secondary ARTs and a core ART, MGG_00230, generated with mTM-align. The structural core is in magnata. **F**. The surface structures of secondary ARTs, MGG_16321 and MGG_09666. The left models indicate the conserved structural core in magnata. The right models display interaction interfaces in red predicted by MaSIF. **G**. A proposed mechanism of the ART evolution in *M. oryzae*.

The other members in this cluster are highly divergent from the core ARTs, and the next largest BLASTP-based homologous group is composed of six members, which we designate as secondary ARTs. The homology from core ARTs to the secondary ARTs is complex (Fig. S17). Often, homology detection is only unidirectional, and shared homology to intermediate sequence (e.g. MGG_09666) is required to infer evolutionary connections (Fig. 5*D*) (70). An explanation for such a complex network is the inclusion of different homologous groups by false homology detection. Alternatively, extreme sequence diversification of the secondary ARTs may also provide adequate rationales.

To understand how the secondary ARTs may evolve, we focused on the four members with predicted structures (Table S5). Upon superpositions of their structures against a core ART, MGG_00230, noticeable differences in evolutionary patterns appeared in sequences and structures. First, the catalytic residues were absent, implying these proteins may be pseudo-enzymes incapable of mediating NAD^+^-dependent ADP-ribosylation (Fig. S18) (71). Second, structural conservation is skewed towards one side of the core ART, although structural similarities do exist between the core and secondary ARTs (Fig. 5*E*). As the conserved region may play a functional role, we employed MaSIF, a geometric deep learning algorithm, to predict the protein interaction interface (72). The comparison of the surface structures revealed that the predicted interface overlaps with the structurally conserved regions, while the overall fingerprints of the paralogs are distinct (Fig. 5*F*). This possibly indicates that the structural core may constitute the backbone of the interface, whereas the other sequence around the core may determine the shape complementarity for substrates or protein targets.

The data suggest a hypothesis in the ART evolution in *M. oryzae*. Canonical, bifunctional ARTs, which bind to possibly essential host targets and transfer an ADP moiety, first emerged and formed the core ART cluster by frequent duplication and diversification (Fig. 5*G*). One of the paralogs lost catalytic residues necessary to metabolize NAD^+^ and began to deviate from the canonical ARTs in evolution. Instead of being selected against, this paralog evolved the remaining protein-protein interaction interface and became specialized as host protein binders. Eventually, additional pseudo-ARTs arose by duplication events, and they subsequently diversified for different host proteins.

## Discussion

### Advances and obstacles in effector structure prediction

Effector structure prediction is intrinsically challenging for template-based modeling. Expectedly, the homology modeling by I-TASSER only resolves a small subset of the secretome. This suggests that secretome-wide structure prediction with homology modeling is not a rational choice to study effector biology. With the advent of deep learning-based, reliable free-modeling, the situation has now changed. TrRosetta modeled a large number of protein structures I-TASSER could not. Even for the commonly predicted structures, the TrRosetta models predominantly displayed higher estimated precision. Such results support that free-modeling with TrRosetta sufficiently captures a large set of effector structures. By supplementing with homology modeling for difficult targets with available templates, such as Avr1-CO39, we can now generate good-quality structures representative of the overall secretome in the pathogen’s genome. That is, secretome-wide structure prediction is now a reasonable decision to unravel the unknown, and diverse downstream analyses that require such resources can be conducted.

Some challenges remain to be resolved. Both TrRosetta and I-TASSER could not generate reliable structures for 559 proteins that are typically small and lack homologs. Most of these proteins failed to retrieve a sufficient number of diverse homologs necessary for co-evolutionary inference, attributed to homology detection failure or recent origins of effectors. In such cases, more genomic data will only increase the number of homologs but unlikely enhance the evolutionary diversity. Predicting their structures with TrRosetta, therefore, heavily relies on the performance of the trained machine learning algorithm without co-evolutionary information, which remains intrinsically difficult. In the CASP 14 in 2020, AlphaFold 2 demonstrated its exceptional success in protein structure prediction (73). Once algorithms with a similar level of performance become available for research, we will be able to not only enhance the quality of the predicted structures but also reveal the structures of the unmodeled proteins in this study.

### Sensitive sequence-based clustering should be conducted for homologous group identification

The lack of homologs may be attributed to the recent origins of effectors. Alternatively, homology detection failure could be another cause as rapid sequence diversification is a common theme in effector evolution. The scenario we proposed for the ARTs may at least partially explain how some effectors follow an unusual evolutionary trajectory and eventually lose detectable homology from their canonical paralogs. Such rapid evolution limits homolog collection and hinders structure prediction. The proteins may end up with poor-quality structural models that cannot be reliably used for structural comparisons and classifications. Nonetheless, distant homology to the rapidly evolving proteins may be still detectable with sensitive homology search methods. Therefore, if homology-based clustering can reveal the membership of these proteins, the structural annotations can be possibly transferred from other members with more reliable structures to infer their putative roles.

Sensitive, sequence-based clustering also assists in resolving ambiguity structure-based inference alone cannot. For instance, MoCDIP1 displays similarity to pectin lyases, yet the highly conserved residues in its structural match are missing. Sequence-based clustering places MoCDIP1 to Cluster 24 with other pectin lyases, leading to the consensus between the two approaches. Similarly, the structural classification assigns an SspB-like fold with low confidence to two members in Cluster 29.

However, the other majority of the homologous members display the Cyanovirin-N fold. Together with the detectable CVNH domain in a few members of this cluster, the consensus indicates that the proteins function for lectin-binding through the CVNH domain and structures. Structure-based annotation could be sensitive to the quality of the predicted structures, structure search method and structure databases; therefore, examining such consensus between sequence and structure-based assignments would be critical prior to downstream analyses.

### Sequence-unrelated, structurally similar effectors are a common theme in effector evolution

MAX effectors represent the sequence-unrelated, structurally similar effectors in *M. oryzae*. Their origin was, however, suggested to be ancient, and duplications as well as diversifying evolution more likely explain the frequent occurrence of these effectors within an organism and across the phytopathogens (16, 74). With protein structure prediction, we also illustrated that putative effectors with the alpha-beta plait topology or the γ-crystallin-like fold may be sequence-unrelated but structurally similar. Additionally, the structural analogy to other phytopathogens’ effectors appeared to be common. This indicates that there could be structural commonalities in phytopathogens that we do not yet fully understand. That is, although the pathogens may utilize effectors seemingly unrelated by sequences, they might have originated from a common ancestor and function similarly with relatively well-maintained ancestral structural folds. Although the lack of effector structures has made it difficult to test such hypotheses, we propose that effecterome-wide structure prediction and structure-incorporated comparative genomic analyses will be able to provide a solution to this problem.

Although structural comparisons successfully identified many analogous effector clusters, the method might have not fully retrieved the cluster members. These effectors would eventually remain as singletons. If investigating sequence-unrelated, structurally similar effectors in *M. oryzae* is of interest, the lower sensitivity of the clustering could be complemented by relaxing the stringency for the structural prediction and clustering. The overall performance of the clustering is yet to be examined. The previous study estimated that *M. oryzae* isolates would encode about 40 to 60 putative MAX effectors, while we identified up to 16 candidates by decreasing the TM-score cut-off by 0.03 in clustering (16). Among the 17 MAX effector candidates from *M. oryzae* 70-15 the authors identified with sensitive PSI-BLAST search, only 8 members were in common. This points to possible issues in sensitivity and precision associated with sequence-based and structure-based MAX effector identification. Once more structures become available, the performance of each methodology will be able to be evaluated.

### The emergence of pseudo-enzymes – can this be a common theme in effector evolution?

We proposed the emergence of pseudo-enzymes and their subsequent diversification in the ART evolution (Fig. 5*G*). Although the discussion about this notion is yet scarce in effector evolution, it is not entirely new. *B. graminis* secretes a significant number of effectors that adopt the RNase fold; however, many of these effectors lack essential residues for catalytic activities, suggesting that they may be pseudo-enzymes (19). A recent study demonstrated that an RNase-like effector without a canonical enzymatic activity still has a functional role in pathogenesis (21). It interacts with not only host RNAs but also a host ribosomal protein, PR10, thereby interfering with host cellular activities.

The expansion of pseudo-enzymes and the validation work of the function raise an intriguing perspective in effector evolution. For both *M. oryzae* pseudo-ARTs and *B. graminis* RNase-like effectors, the ancestral, canonical protein would likely have properties to bind to important host targets. Loss of catalytic activities would not be evolutionarily deleterious to the effector, if its binding to the host targets was sufficient for pathogenesis. Instead, this event would relax evolutionary constraints to maintain the enzymatic functions in both sequence and structure, eventually opening new paths in evolutionary trajectories that canonical ARTs or RNAse-like effectors could not access. Therefore, the ancestor could become specialized as a binder, and frequent duplication and diversification subsequently allowed the paralogous proteins to follow other accessible evolutionary paths, eventually leading to an expanded protein family that targets a diverse set of host molecules.

### Predicted structures alone cannot tell the entire story of effector biology

Although well predicted structural information is valuable, it alone cannot tell the entire story of effector biology. The structure-based functional inference is analogous to sequence-based functional annotation transfer. Both require not only the predicted structures or sequences of good quality but also the existence of references and their known functions. The scope of solved effector structures and our understanding of effector functions are currently limited. Even with predicted structures of good quality, we may not be able to derive more information from them. Molecular and structural biology work remains critical, regardless of the improvement in structure prediction algorithms. Our work provides a unified structural genomics resource that can be used to group and prioritize candidate effectors for further analyses.

Genome-scale protein structure prediction is still time-consuming and computationally intensive. However, with the advances in machine learning and parallel computing, the field of protein structure prediction is rapidly evolving to challenge this problem. The growing structural data will shift our perspectives on evolution towards the three-dimensional space, unrestricted to primary sequences. We also foresee that structural genomics will be applied to not only the effectors of phytopathogens but also much larger proteomes of their hosts. Such datasets will facilitate understanding the interplay between effectors and host proteins and co-evolution stemming from this interaction.

## Materials and Methods

### Secretome prediction and benchmarking data collection

The 12,755 protein sequences of *M. oryzae* strain 70-15 were downloaded from Ensembl (75). We utilized the neural network of SignalP v3.0, one of the most sensitive to defect fungal effectors, to identify putative secreted proteins (76, 77). SignalP initially predicted 2,348 secreted proteins with the D-score ≥ 0.43. 119 possible false positives were removed as their predicted signal peptides overlapped with PFAM domains annotated with InterProScan v5.30-69.0 over ≥ 10 amino acids (78). Following the predicted cleavage site based on the Y-score from SignalP, mature protein sequences were generated. We used TMHMM v2.0 to eliminate 344 putative integral membrane proteins with one or more transmembrane helices in the mature proteins (79). Lastly, 25 mature proteins ≥ 1000 amino acids were removed, and only the longest isoform was selected. In total, 1,854 proteins remained for the analysis. For benchmarking, we collected 24 representative *Magnaporthe* proteins with solved structures from the PDB (27). The full-length sequences were identified for PDB entries with BLAST against the nr database in the National Center of Biotechnology Center (NCBI) (80). The protein was classified as secreted if its full-length sequences satisfied the above criteria.

### Gene prediction for the Magnaporthales

We obtained 248 genome assemblies for the organisms in the order Magnaporthales from the NCBI. As the primary purpose of the genome annotation was to collect homologs of the predicted secreted proteins of *M. oryzae*, we mainly relied on Liftoff v1.3.0 to transfer existing annotations of *M. oryzae* 70-15 to a target genome (81). We then utilized BRAKER v2.1.5 to predict additional gene models uncaptured by Liftoff (82–88). For 222 *M. oryzae* genomes, for instance, the reference annotation of *M. oryzae* 70-15 was initially mapped into each target genome with Minimap v2.17-r974-dirty (89). The gene models with ≥ 98% coverage and ≥ 40% sequence identity were annotated by Liftoff and collected if no premature stop codon existed (-a 0.98 -s 0.4 -sc 0.4 -copies -exclude_partial). Prior to running BRAKER, the repeat library for the reference genome was generated by RepeatModeler v2.0.1 (90). RepeatMasker v4.1.0 was used to soft-mask each target genome (91). The fungal proteomes collected from OrthoDB v10 and the genome annotations generated by the previous study and available in the NCBI for the order Maganaporthales were used as protein evidence for BRAKER (14, 92). Among the final annotations, we selected all the gene models not overlapping with the ones previously predicted by Liftoff. For other species in Magnaporthales, we used the annotation sets of their closely related species as the reference and followed the annotation pipeline.

### Generation of a deep multiple sequence alignment

To search for homologs of a target protein, DeepMSA was used (93). DeepMSA iteratively searches the Uniclust30, Uniref90 and Metaclust databases to construct a MSA (94–96). We supplemented the Metaclust database with additional fungal protein sequences to facilitate collecting diverse fungal homologs. The fungal datasets consisted of 1689 annotated proteomes from the Joint Genome Institute and the 249 Magnaporthales annotations we produced (97). The two datasets were concatenated, and gene models with premature stop codons were removed. The 25,077,589 gene models were clustered with Linclust from MMseqs2 to reduce redundancy (--min-seq-id 0.95 --cov-mode 1 -c 0.99) (98, 99). The resulting 17,679,966 representative gene models were appended to the Metaclust database. DeepMSA was run on these databases to generate a MSA for each secreted protein without the final filtering step.

### Protein structure modeling and final structure selection

The protein structures of all the candidate secreted proteins were predicted by two different methods: homology modeling by I-TASSER v5.1 and free modeling by TrRosetta (25, 26). For I-TASSER, the template library was downloaded from the I-TASSER server (08/24/2020). I-TASSER was run in the light mode with the MSA from the previous step converted into the PSI-BLAST-readable format using the a3m2mtx script in the DeepMSA package (-a3m -neff 7). For the benchmarking set, any homologous templates with ≥ 70% sequence identity were excluded (-homoflag benchmark -idcut 0.7). We used the estimated TM-score of the first model as a measure of precision, and the predicted structures with the mean TM-score > 0.5 were considered reliable.

For TrRosetta, we selectively filtered the MSAs, limiting their sizes to 6,000 sequences to prevent unnecessarily deep, large MSAs. If the MSA from deepMSA was larger than the limit, sequences with ≥ 75% of the query coverage were first sampled from high to low sequence identity over the aligned regions. If necessary, sequences with ≥ 50% of the query coverage were additionally sampled. In the sampling, only the sequences with identity < 90% were counted. TrRosetta was run with the filtered MSAs to predict inter-residue orientations and distances, and we generated five full-atom models with PyRosetta (100). The model with the lowest energy score was chosen as a final model. We used the average probability of the top L predicted long+medium range contacts (|i-j|>12) estimated by the top_prob.py script in the TrRosetta suite as precision measurement. A structure with the probability > 0.5 was considered reliable. Among the outputs from I-TASSER and TrRosetta, the structure with higher estimated precision was selected as a final model. We used Pymol v2.4.0 to visualize the structures (101).

For the benchmarking data, the actual precision of the predicted structures was accessed by superposing the computational model against the experimental one with TM-align (28). The TM-score normalized for a smaller protein was chosen.

### Functional and structural annotations of the secretome

We employed InterProScan v5.30-69.0 to identify homologous PFAM v31.0 entries (102). PaperBLAST and PHI-base were used to search for *M. oryzae* effectors and their homologs in other pathogens (103, 104). The secondary structures and disordered residues were predicted with PSIPRED v4.0 and DISOPRED v3.16 (105, 106). The predicted structures were compared with RUPEE against the SCOPe v2.07 and CATH v4.2.0 databases as well as the PDB chains (TOP_ALIGNED, FULL_LENGTH) (27, 34–36). Regardless of the confidence scores, a top hit was reported.

### Network analysis using homology and structures

We identified homology using three different methods: sequence-to-sequence search with BLASTP v2.7.1, profile-to-sequence search with HHblits v3.1.0, and profile-to-profile search with HHsearch v3.1.0 (48, 49). The MSAs generated in the previous step represented the profiles. In all cases, only the pairs with E-value < 10^−4^ and bidirectional coverage > 50% were regarded significant. The structural similarities were compared with TM-align by superposing each pair of structures predicted with the estimated precision > 0.5 by TrRosetta or > 0.55 by I-TASSER. We increased the cut-off for I-TASSER as the structures with the estimated precision around 0.5 introduced spurious clustering, and the sources of their selected templates were often distantly related organisms like humans. We also did not use any proteins with ≥ 35% disordered residues for structural comparisons, as these proteins tended to be modeled by I-TASSER with similar homologous templates and thus introduce biases in the similarity search. The two compared structures were considered similar if the TM-score was higher than 0.5 for both structures or 0.6 and 0.4 for each. We used the igraph package v1.2.4.1 in R v3.6.1 for the network analyses and to reveal the cluster membership of secreted proteins (107, 108)

## Acknowledgments

We appreciate Brian Staskawicz for allowing us to utilize the computational resources and Jianyi Yang for his advice on structure prediction. We also thank China Shaw, Daniil Prigozhin and Pierre Joubert for the critical review of the manuscript. This research relied on the Savio computational cluster resource provided by the Berkeley Research Computing program at the University of California, Berkeley. Kyungyong Seong is supported by Berkeley Fellowship.

## Data availability

The genome annotations, MSAs and structures generated for the secreted proteins, and the data used for network analyses are deposited in Zenodo (10.5281/zenodo.4456015).

